# Clustering and erratic movement patterns of syringe-injected versus mosquito-inoculated malaria sporozoites underlie decreased infectivity

**DOI:** 10.1101/2020.10.21.348573

**Authors:** C.M. de Korne, B.M.F. Winkel, M.N. van Oosterom, S. Chevalley-Maurel, H.M. Houwing, J.C. Sijtsma, E. Baalbergen, B.M.D. Franke-Fayard, F.W.B. van Leeuwen, M. Roestenberg

**Affiliations:** Department of Infectious Diseases and Parasitology, Leiden University Medical Center, Albinusdreef 2, 2333 ZA Leiden, The Netherlands; Interventional Molecular Imaging laboratory, Department of Radiology, Leiden University Medical Center, Albinusdreef 2, 2333 ZA Leiden, The Netherlands

## Abstract

Live attenuated malaria sporozoites are promising vaccine candidates, however, their efficacy critically depends on their capability to reach and infect the host liver. Administration via mosquito inoculation is by far the most potent method for inducing immunity, but highly unpractical. Here, we observed that intradermal syringe-injected *Plasmodium berghei* sporozoites (^syr^SPZ) were three-fold less efficient in migrating to and infecting mouse liver compared to mosquito-inoculated sporozoites (^msq^SPZ). This was related to a clustered dermal distribution (2-fold decreased median distance between ^syr^SPZ vs ^msq^SPZ) and, more importantly, a 1.4-fold significantly slower and more erratic movement pattern. These erratic movement patterns were likely caused by alteration of dermal tissue morphology (>15 μm intercellular gaps) due to injection pressure and may critically decrease sporozoite infectivity. These results suggest that novel microvolume-based administration technologies hold promise for replicating the success of mosquito-inoculated live attenuated sporozoite vaccines.

## INTRODUCTION

Since 2014, the number of cases of malaria worldwide has remained at 200 million annually leading to more than 400 thousand deaths every year, with children in sub-Saharan Africa bearing the greatest burden (according to the World Health Organization, World Malaria Report 2019). This high morbidity and mortality underlines the pressing need for an effective vaccine to support control programs. Live attenuated *Plasmodium falciparum (Pf)* sporozoites are promising malaria vaccine candidates which currently are in clinical development (PfSPZ Vaccine, PfSPZ CVac, PfSPZ-GA1(1)) and have the potential of inducing up to 100% sterile immunity(2, 3).

To make live attenuated sporozoites amenable for large scale immunization, the US-based biotech Sanaria has developed tools to isolate, purify and cryopreserve sporozoites for injection. Unfortunately, the dermal or subcutaneous injection of sporozoites provided suboptimal protective efficacy(4). Although much better efficacy was obtained when injecting high numbers of sporozoites intravenously(5), the intradermal, intramuscular or subcutaneous administration routes of low numbers of sporozoites are preferred to facilitate global administration to infants in endemic countries at a low cost-of-goods. A better understanding of the differences between the potent mosquito-inoculated sporozoites and the unsuccessful syringe-injected sporozoites is needed to boost the development of practical and efficacious attenuated sporozoite vaccines.

The potency of attenuated sporozoite vaccines critically depends on the ability of the sporozoite to migrate to and infect the host liver. Transgenic luciferase-expressing sporozoites and bioluminescence-based visualisation of parasites in mice provide a macroscopic imaging platform to study the liver stage parasite burden after different routes of administration(6). These studies indicated that mosquito-inoculated sporozoites migrate to the liver much more efficiently as compared to intradermally injected parasites(7–10). The subsequent development of fluorophore-expressing sporozoites and a sporozoite fluorescent labelling approach has allowed for more detailed microscopic studies on the motility of individual sporozoites, both *in vitro* and in skin(11, 12). Moreover, automated analysis of sporozoite motility now provides a platform to quantitatively study sporozoite motility under different conditions(13–17).

We now aimed to unravel the factors underlying the difference in potency and infectivity between mosquito-inoculated and intradermal syringe-injected sporozoites (Scheme 1). For this we microscopically examined the dermal site, quantitatively assessed the distribution of sporozoites and their motility patterns after inoculation by mosquito bite (^msq^SPZ) and injection by needle and syringe (^syr^SPZ) through automated image analysis. We assessed liver stage parasite burden through bioluminescence assays.

**Scheme 1.**
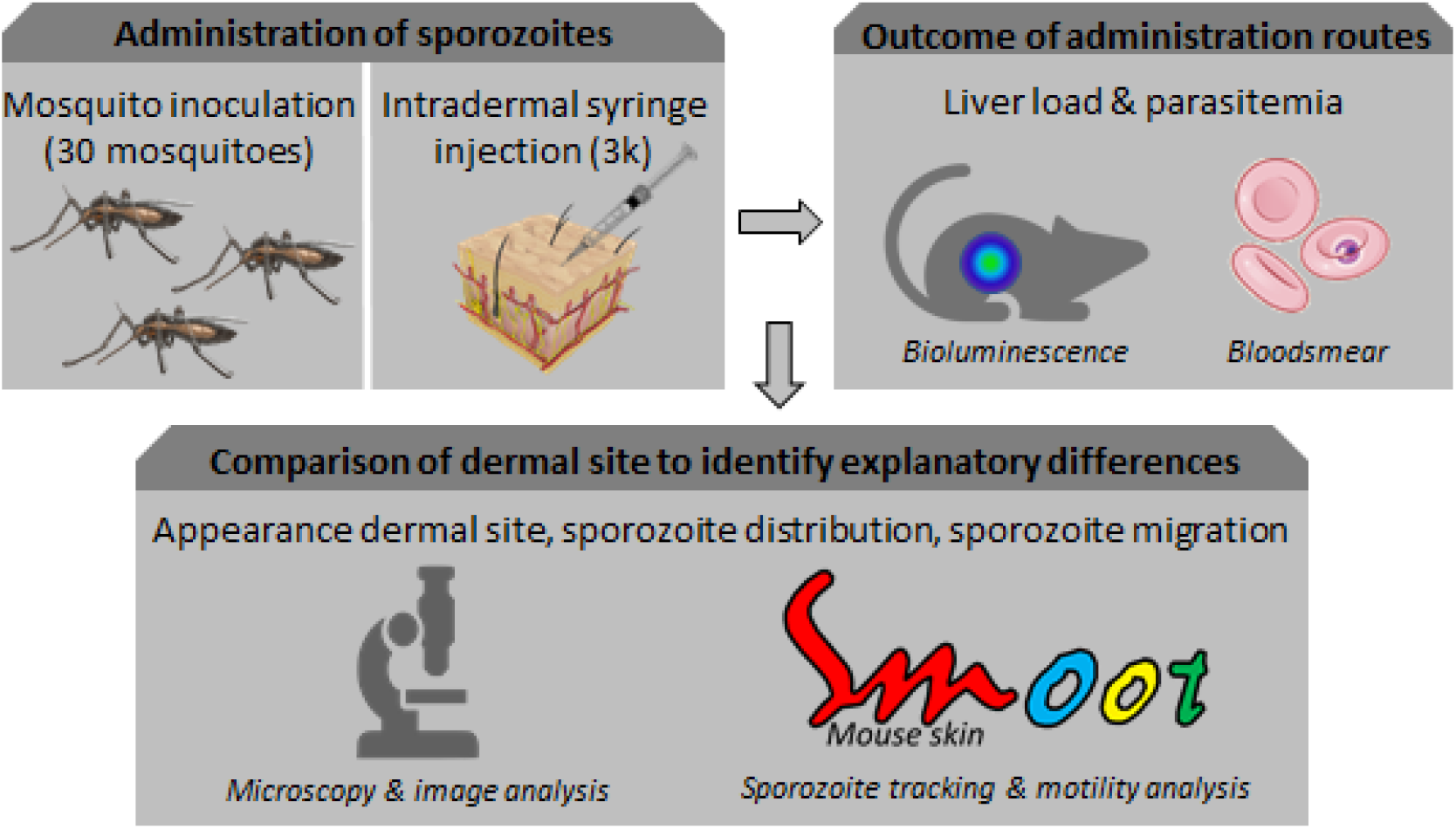
Study design. Sporozoites were administered via mosquito inoculation or intradermal syringe injection. Liver load was subsequently assessed by bioluminescence and blood smear patency. Detailed analysis of 1) the appearance of the dermal site, 2) the sporozoite distribution and 3) sporozoite migration behaviour was performed to reveal underlying mechanisms of decreased infectivity.

## METHODS

### Rodent experiments

Mouse experiments were performed with Female Swiss OF1 mice (6-7 weeks old; Charles River. All animal experiments were granted a licence by the Competent Authority after an advice on the ethical evaluation by the Animal Experiments Committee Leiden (AVD1160020173304). All experiments were performed in accordance with the Experiments on Animals Act (Wod, 2014), the applicable legislation in the Netherlands and in accordance with the European guidelines (EU directive no. 2010/63/EU) regarding the protection of animals used for scientific purposes. All experiments were performed in a licenced establishment for the use of experimental animals (LUMC). Mice were housed in individually ventilated cages furnished with autoclaved aspen woodchip, fun tunnel, wood chew block and nestlets at 21 ± 2°C under a 12:12 hr light-dark cycle at a relative humidity of 55 ± 10%.

### Sporozoite production

Naive mice were infected with the rodent malaria species *P. berghei* as described previously(7). The transgenic line 1868cl1 expressing mCherry and luciferase under the constitutive HSP70 and eef1a promotors respectively (*Pb*ANKA-mCherry_hsp70_+Luc_eef1α_; line RMgm-1320, www.pberghei.eu) was used. The infected mice were anesthetized and *Anopheles stephensi* female mosquitoes were infected by feeding on the gametocytaemic mice as described previously(18). The mosquitoes were kept at a temperature of 21 °C and 80% humidity until use.

### Sporozoite administration

After anesthetizing mice, *P. berghei* 1868cl1 sporozoites were administered using two different methods:

1. ^msq^SPZ were delivered by shaving the mice and 1 cm^2^ skin was exposed for 15 minutes to around 30 infected mosquitoes (exact numbers are specified per experiment). Blood-fed mosquitoes were counted and the presence of sporozoites in their salivary glands was confirmed using the quantitative analysis of the luciferase activity after putting the mosquitoes in a 20 μl drop D-luciferin (8 mg/ml in phosphate-buffered saline (PBS)).
2. ^syr^SPZ were obtained by manual dissection of the salivary glands of infected female *Anopheles stephensi* mosquitoes 20-24 days post infection. The salivary glands were collected and homogenized to release sporozoites in Roswell Park Memorial Institute medium (RPMI; Thermo Fisher Scientific) enriched with 10% fetal bovine serum (Life Technologies Inc.), unless otherwise stated. The free sporozoites were counted in a Bürker counting chamber using phase-contrast microscopy to prepare the injection samples. Directly, after dissection and counting, 10 μl sample containing 3000 sporozoites was administered by injection into the skin of the upper thigh using an insulin syringe (Becton Dickinson, Micro-Fine+, 0.5 ml; 0.30 × 8.0 mm, 30G). The number of sporozoites delivered by exposure to 30 infected mosquitoes was considered consistent with intradermal delivery of 3000 sporozoites via syringe injection based on available literature (mean number of sporozoites inoculated per mosquito: 116 ± 28(19)).

### Quantification of parasite liver load and prepatent period

The liver stage of the *P. berghei* infection in 9 mice (6 mice challenged by 33 (IQR: 30-33) mosquito bites of which 25 (IQR: 22-28) contained blood and 3 mice challenged by syringe injection of sporozoites in RPMI) was visualized and quantified by measuring the luciferase activity in the liver at 44 hours after the challenge with sporozoites using the IVIS Lumina II Imaging System (Perkin Elmer Life Sciences). Before imaging, the mice were shaved and anesthetized. IVIS measurements were performed within 8 minutes after subcutaneously injection of D-luciferin in the neck (100 mg/kg in PBS; Caliper Life Sciences). Image analysis was performed using the Living Image^®^ 4.4 software (Perkin Elmer Life Sciences). Infected mice were monitored for blood-stage infections by Giemsa-stained blood smear until day 9 post infection. The prepatent period (measured in days after sporozoite challenge) was defined as the first day at which blood stage infection with a parasitaemia of >0.5% was observed.

### Quantification of sporozoites by PCR

Directly after sporozoite delivery by 32 (IQR: 32-33) mosquito bites of which 18 (IQR: 17-19) contained blood, the skin of 4 exposed mice was cut out, snap-frozen and stored at −20 °C until further use. Parasite burden was measured by quantitative real time reverse transcription polymerase chain reactions (qRT-PCR). The DNA was extracted from the frozen skin using the QIAamp DNA Micro Kit (Qiagen) following the manufacturer’s instruction. Amplification reactions of each DNA sample were performed in PCR plates (hard-shell PCR plate, #HSP9645; Bio-Rad), in a volume of 25 μl containing 12.5 μl PCR buffer (HotstarTaq mastermix; Qiagen), 0.5 μl MgCl_2_ (25mM), Plasmodium-specific forward and reverse primer (12.5 pmol; Plas-7F 5’-GTTAAGGGAGTGAAGACGATCAGA-3’ and Plas-171R 5’-AACCCAAAGACTTTGATTTCTCATAA-3’; Sigma-Aldrich), PhHV-specific forward and reverse primer (15 pmol; PhHV-267S 5’-GGGCGAATCACAGATTGAATC −3’ and PhHV-337AS 5’-GCGGTTCCAAACGTACCAA −3’; Biolegio), Plasmodium-specific FAM10 labelled detection probe (2.5 pmol; PP FAM 5’-ACCGTCGTAATCTTAACC-3’; Biolegio), PhHV-specific Cy5 double-labelled detection probe (1.25 pmol; PhHV-305TQ Cy5 5’-TTTTTATGTGTCCGCCACCATCTGGATC-3’-BHQ2; Biolegio) and 5 μl of the DNA sample (dilution factor: 10x). Amplification consisted of 15 min at 95°C followed by 50 cycles of 15 s at 95°C, 30 s at 60°C, and 30 s at 72°C. Amplification, detection, and analysis were performed with the CFX96TM real time PCR detection system (Bio-Rad). A calibration curve to assess the sporozoite numbers in the mosquito inoculation samples was generated by analysing skin samples injected with a dilution range of sporozoites (2-step dilution, start: 20.000 sporozoites, 10 samples, n=3 performed in duplo, Sup. Figure 1).

**Figure 1.**
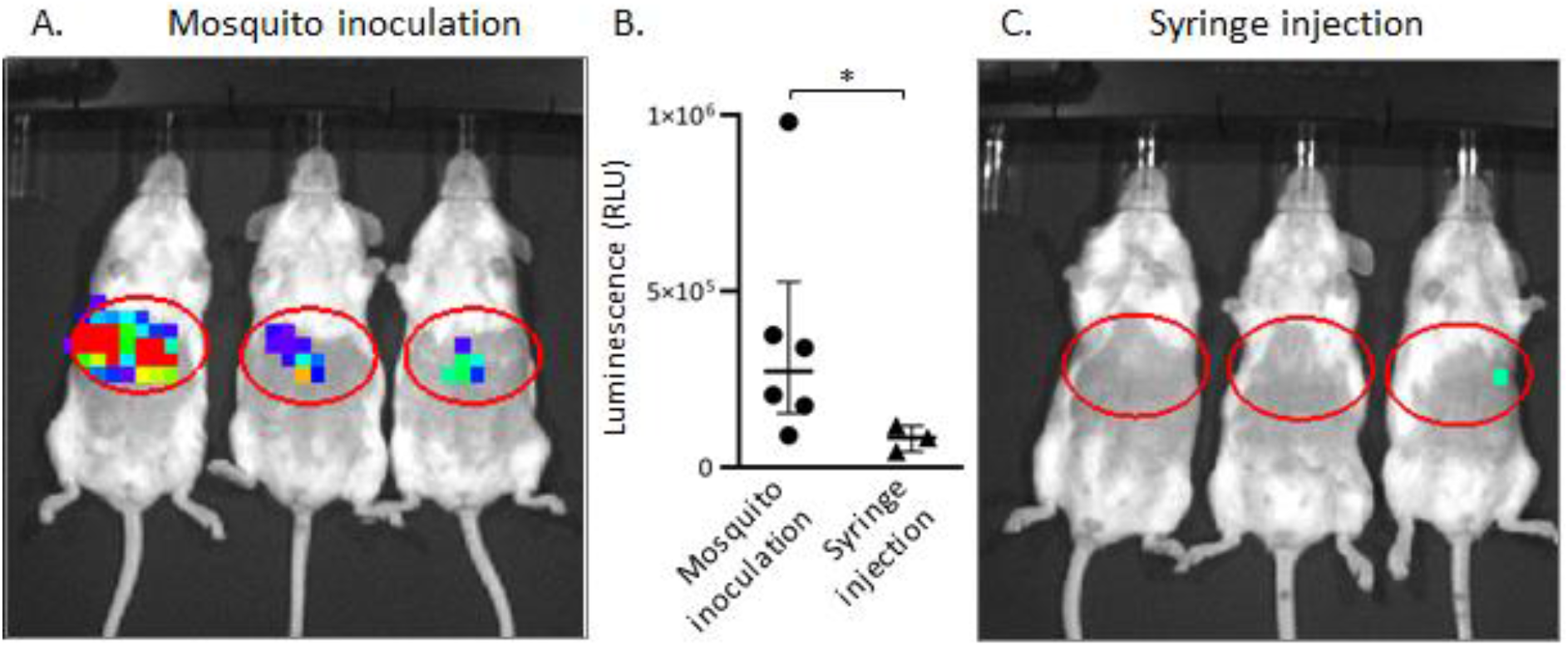
Outcome different administration routes. (A, C) In vivo images which show the liver load 44 hours post infection via 33 (IQR: 30-33) mosquito bites (A) or intradermal syringe injection of 3000 sporozoites (C). (B) Average of the luciferase activity in the liver 44 hours after challenge by mosquito inoculation (median: 2.7*10^5^ RLU, IQR: 1.8*10^5^-3.7*10^5^) or intradermal syringe injection (median: 8.5*10^4^ RLU, IQR: 6.5*10^4^-1.0*10^5^)(*p=0.048; Mann-Whitney U test).

### Ex vivo (fluorescent) imaging of the dermal site

Directly after sporozoite delivery, the exposed skin of 8 mice (4 ^msq^SPZ and 4 ^syr^SPZ samples) was excised, covered with a cover slip and imaged using an Andor Dragonfly 500 spinning disk confocal on a Leica DMi8 microscope (Oxford Instruments) or a Leica True Confocal Scanning SP8 microscope (Leica Microsystems). The mCherry expressed by sporozoites was excited with the 561nm laser. A 20x objective (HC PL APO 20x/0.75 IMM CORR CS2) was used, resulting in images of 617 × 617 μm. The experiments were performed at room temperature.

To create overview images of the dermal site, up to 570 fields of views were stitched using the Andor imaging software Fusion (Oxford Instruments). Z-slices were imaged covering an average total depth of 124 μm (IQR: 103-131). Using the Fiji package for the open source software ImageJ(20), the three-dimensional z-stack was reduced into a two-dimensional image using maximum intensity projection (retrieves the level of maximum intensity along the z axis for each x,y position) and was converted into a binary image only showing the sporozoites. This image was further processed in two different ways: 1) The Gaussian blur filter was applied and a pseudo colour image was created by applying the inferno colour lookup table in Fiji to visualize the location and density of the sporozoites, 2) The coordinates of the individual sporozoites in the ^msq^SPZ and ^syr^SPZ samples were determined and the nearest neighbour distance (distance between the centre points of neighbouring sporozoites) was calculated using the ImageJ Nnd plugin. The number of sporozoites residing within or in close proximity to hematomas was determined using circular ROIs with a diameter of 650 μm and around the centre of the hematoma. The shape of the cells visible at the zoom-in brightfield images (after mosquito inoculation n=164, after syringe injection n=203) was described by two shape descriptors available within ImageJ, namely roundness: 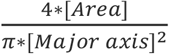 with a value of 1.0 indicating a perfect circle and the Feret’s diameter: the longest distance between any two points along the cell membrane.

### Sporozoite motility

To analyse sporozoite motility, movies were recorded with a frame rate of 35-40 frames per minute, 200 frames per movie. Recording ^msq^SPZ samples (n=6) yielded a total of 3400 frames. Recording ^syr^SPZ samples (n=8) yielded a total of 3600 frames. Maximum intensity projections of the recorded microscopy movies were generated using ImageJ. The motility of the sporozoites was analysed using SMOOT_mouse skin_, an in-house developed software program, written in the MATLAB programming environment (MathWorks). This tool is an adapted version of the SMOOT_human skin_(12, 17) and SMOOT_*In vitro*_(14) tools previously described. Via SMOOT_mouse skin_, the sporozoites could be segmented per movie frame, based on their fluorescence signal intensity, size and crescent shape. The median pixel locations of the segmented sporozoites were connected into full sporozoite tracks.

First, the sporozoite tracks were characterized as motile or stationary based on their displacement, using a displacement cut-off of 21 pixels which corresponds to the length of a sporozoite. Subsequently, the tracks of the motile sporozoites were subdivided into defined movement patterns: sharp turn, slight turn and linear segments. Third, for the motile sporozoites the mean squared displacement at frame level, the average velocity of their tracks and the nearest neighbour distance per track was calculated and the tortuosity of the tracks was described via the straightness index (the ratio between the total length of the direct path between the start and the end of a track and the total length of the travelled path) and the angular dispersion (the deviation from the mean angle of the track). Finally, the interplay between the angular dispersion, straightness index, velocity and nearest neighbour distance was analysed, for which the number of ^msq^SPZ and ^syr^SPZ tracks within the dataset was equated using random sampling. Twelve subsets of sporozoite tracks were defined based on the angular dispersion (<0.5 and >0.5), straightness index (<0.5 and >0.5) and velocity, (<1 μm/s, 1-2 μm/s, >2 μm/s).

### Statistical analysis

The average and variability of the data was summarised using the mean and standard deviation (SD) for parametric data or the median and interquartile range (IQR) for nonparametric data. For the comparison of groups, the difference between means or medians was assessed using respectively the independent sample T test and the Mann-Whitney U test. For the comparison of a group with a set value the one sample T test was used. For the comparison of the distribution of categorical data the Chi-squared test was used, including a post hoc analysis based on residuals. A univariate general linear model was used to examine the relationship between a continuous and categorical variable. p-values < 0.05 were considered significant, in case of multiple testing the Bonferroni correction was applied to adjust the p-value. All statistical tests were performed by SPSS Statistics (IBM Nederland BV). To compare the velocity distributions of both groups, the distribution was described performing Expectation-Maximization based fitting. The probability density function that could describe the sporozoite velocity distribution consisted of a mixture of 2 normal distributions. The package mixR(21) within the open source R environment(22) was used to define the distribution parameters yielding the best fit.

## RESULTS

### Infectivity of ^msq^SPZ and ^syr^SPZ

At roughly equal numbers of administered ^msq^SPZ and ^syr^SPZ, the parasite liver loads of infected mice, assessed by bioluminescence imaging, were 3.2-fold higher in ^msq^SPZ mice (median: 2.7*10^5^ RLU, IQR: 1.8*10^5^-3.7*10^5^) as compared to ^syr^SPZ mice (median: 8.5*10^4^ RLU, IQR: 6.5*10^4^-1.0*10^5^; p: 0.048; Mann-Whitney U test) (Fig. 1). These results were in line with previous reports(7). The prepatent period of blood stage infection was comparable after infection by mosquito bites or syringe injection (on average 7 days in both groups).

Attempts to quantify the number of ^msq^SPZ by qRT-PCR (Sup. Fig. 1) showed high variability in estimates (median: 6060, range 2203-13,481), which was at least partly caused by technical variability inherent to DNA extraction from skin lysis samples. The estimated average number of ^msq^SPZ delivered did not significantly differ from the targeted 3000 (p: 0.205; one-sample t test).

### Dermal site appearance

The dermal ^msq^SPZ and ^syr^SPZ sites were imaged over an average total depth of 124 μm (IQR: 103-131) by confocal microscopy in order to visualize the sporozoite distribution. In general, ^msq^SPZ were distributed both individually and in clusters dispersed throughout the dermal tissue (Fig. 2A). We found that ^msq^SPZ dermal tissue contained multiple hematomas (median number per sample: 6.5). Interestingly we also found hematomas and ^msq^SPZ in the mouse peritoneum (Sup. Fig. 1 & 2). Of the ^msq^SPZ identified, 9% was found within or in close proximity to the hematomas (example is shown in Fig. 2Aiv), which represented roughly 23% of the hematomas (6/26).

**Figure 2.**
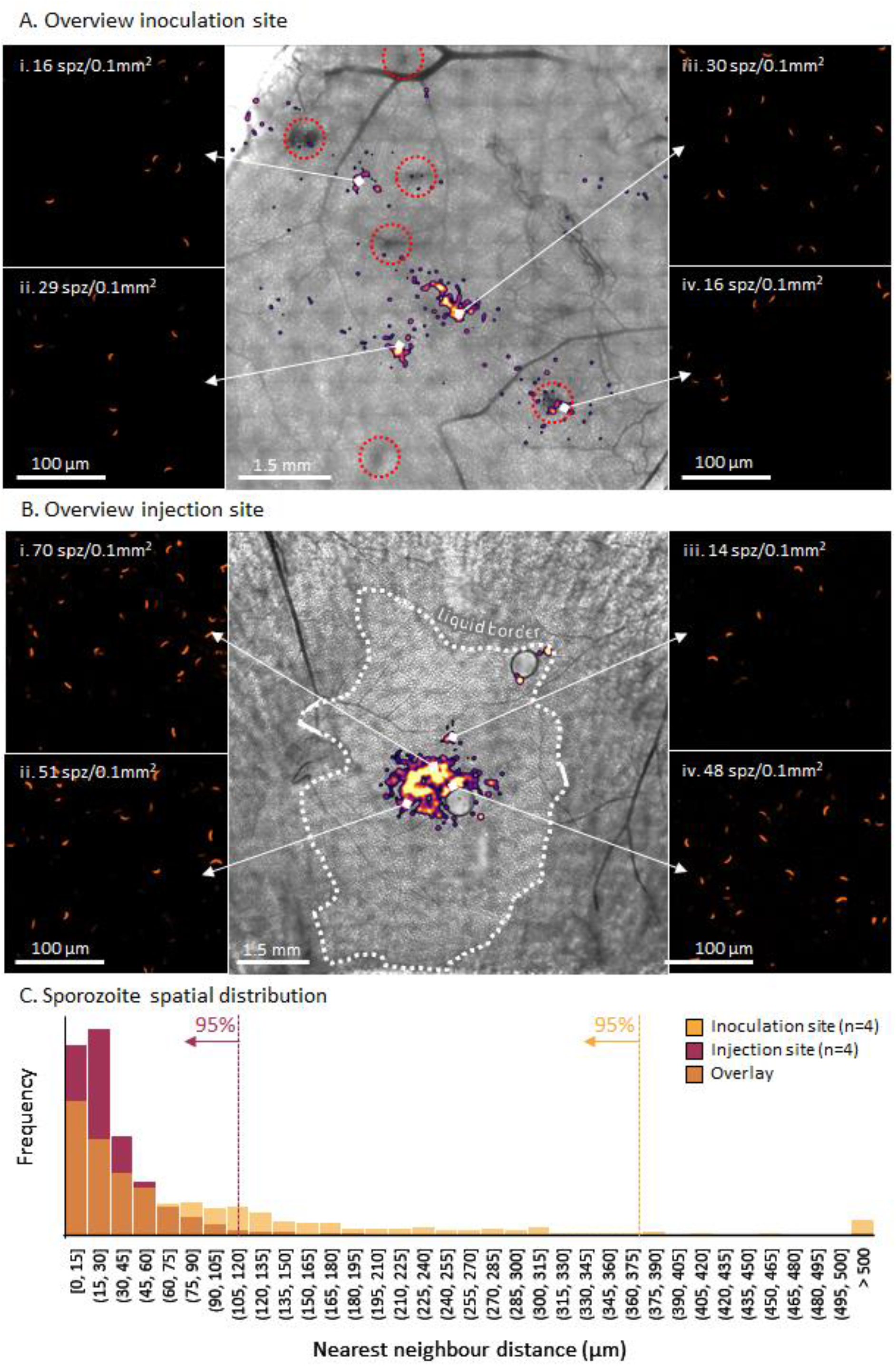
Overview dermal site appearance. (A-B) Overview of the inoculation site after sporozoite delivery by mosquito (A) and the injection site after sporozoite delivery by intradermal syringe injection (B), shown as an overlay of a brightfield image and the sporozoite distribution (pseudo-coloured, blurred fluorescent image) accompanied by zoom-in images showing the individual sporozoites (i-iv). (C) Plot of the nearest neighbour distance for ^msq^SPZ (yellow, median: 55 μm, IQR: 18-132) or ^syr^SPZ (purple, median: 23 μm, IQR: 13-43), the overlap of both distributions is plotted in orange.

In contrast, dermal sites containing ^syr^SPZ showed the injected medium diffused throughout skin, with a single cluster of ^syr^SPZ in the centre of this injection site (Fig. 2B). We did not find hematomas in the ^syr^SPZ dermal tissue, nor ^syr^SPZ in the peritoneum.

The sporozoite distribution was quantified according to their nearest neighbour distance (NND) confirming the dispersed nature of ^msq^SPZ with a median NND of 55 μm (IQR: 18-132), 5% of the ^msq^SPZ were further than 376 μm apart (Fig. 1C). In contrast, ^syr^SPZ were clustered at a median NND of 23 μm (IQR: 13-43), 5% of the ^syr^SPZ were further than 112 μm apart.

Zooming in on the morphology of the skin tissue, we found that after mosquito inoculation, cells remained densely packed resulting in polygonal shaped cells (mean roundness: 0.75 ± 0.11; Feret’s diameter: 82 ± 13 μm) (Fig. 3AB). Conversely, after the syringe injection the interstitial space between the cells was enlarged, leading to >15 μm gaps between cells and a change in cell shape towards significantly more rounded cells (mean roundness: 0.87 ± 0.07, p<0.001, independent sample T test; Feret’s diameter: 68 ± 10 μm, p<0.001, independent sample T test) (Fig. 3AB). The projection of ^msq^SPZ and ^syr^SPZ tracks on top of brightfield images showing tissue morphology revealed that the altered tissue morphology was accompanied by altered movement patterns (Fig. 3C).

**Figure 3.**
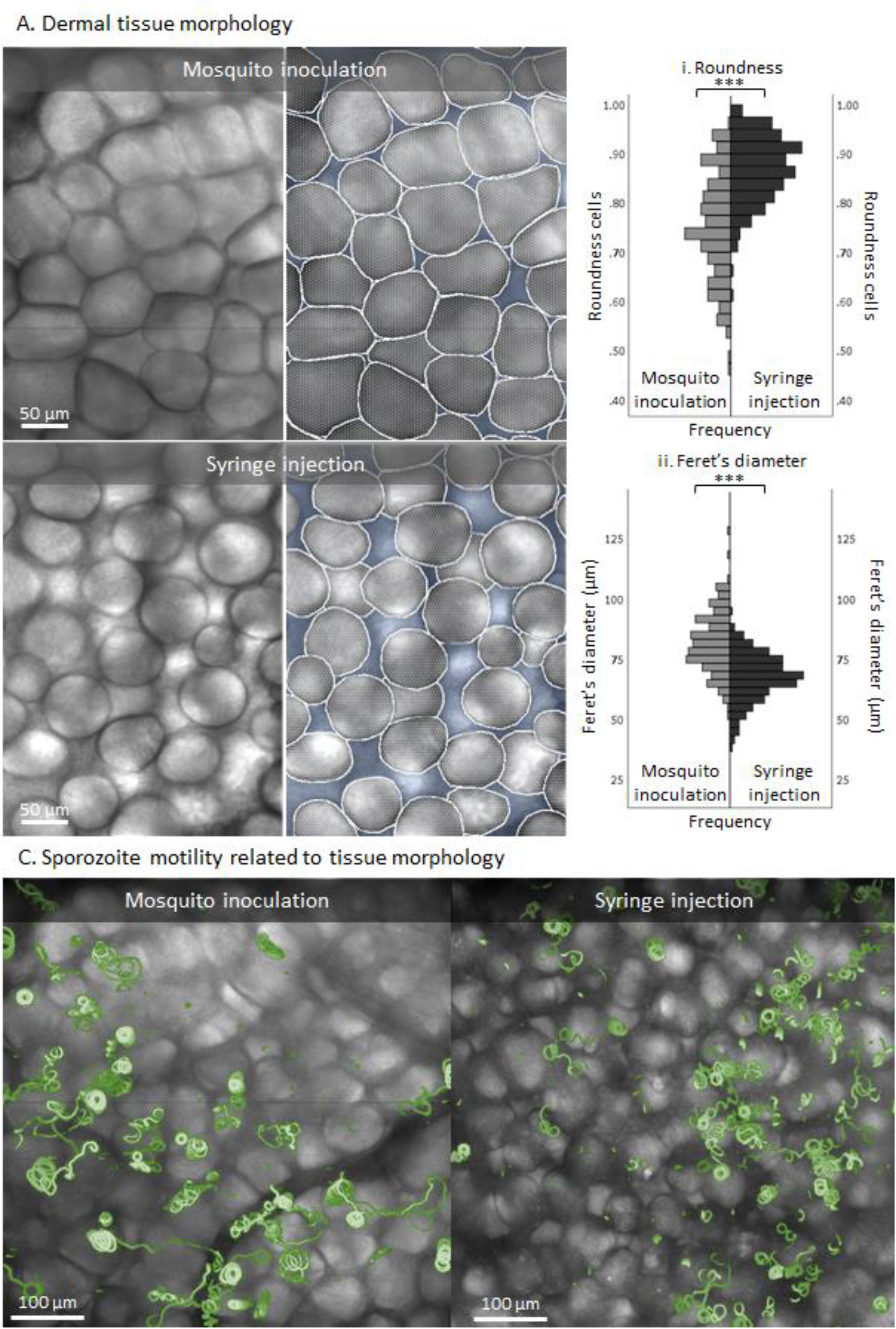
Zoom-in on dermal tissue morphology after sporozoite delivery. (A) Zoom-in on the tissue morphology of the inoculation site after sporozoite delivery by mosquito and the injection site after sporozoite delivery by intradermal syringe injection. Based on the brightfield images, the cells (depicted in white) and the interstitial space (depicted in blue) were segmented. (B) Quantification of the cell shapes found after mosquito inoculation (n=164) and syringe injection (n=203) using roundness (i) and the Feret’s diameter (the longest distance between any two points along the cell membrane) (ii) as measures. ***p<0.001; independent sample T test. (C) Overview of the dermal site shown as an overlay of a brightfield image and a map of mosquito-inoculated and syringe-injected sporozoite tracks (depicted in green).

### Intradermal sporozoite motility

#### Directionality

After delivery, the majority of sporozoites displayed tortuous movement through the dermal tissue, representative examples are shown in Figure 4A. In total, the movement of 566 ^msq^SPZ and 1079 ^syr^SPZ could be captured and analysed. Both ^msq^SPZ and ^syr^SPZ were equally motile (respectively 89% and 88%).

**Figure 4.**
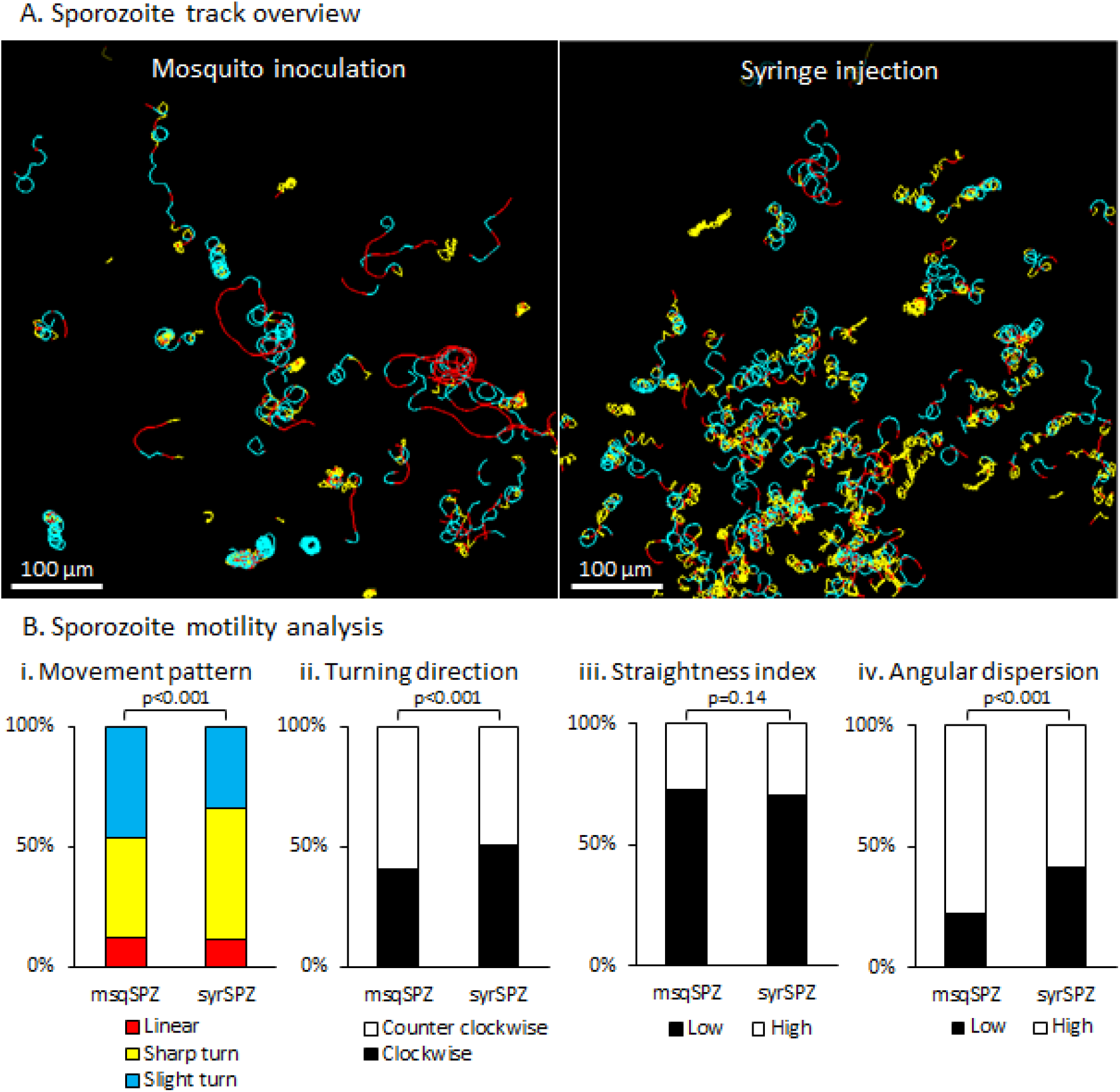
Movement pattern of sporozoites. (A) Overview of the inoculation site after sporozoite delivery by mosquito and the injection site after sporozoite delivery by intradermal syringe injection, shown as a map of sporozoite tracks, colour-coded for movement pattern (sharp turns in yellow, slight turns in blue, linear movement in red). (B) Quantification of the different aspects of sporozoite motility after administration by mosquito or syringe: (i) The movement pattern distribution based at frames, (ii) the percentage of clockwise and counter clockwise segments (iii) the straightness index of the tracks (low: <0.5, high: >0.5) and (iv) the angular dispersion of the tracks (low: <0.5, high: >0.5). p-values obtained by Chi-square test.

Following both administration routes, the tracks of the sporozoites were highly curved. Traditional motility measures such as the mean squared displacement were thus unsuitable to accurately describe the migration behaviour of both groups of sporozoites (Sup. Fig. 3). Therefore, we included other parameters to investigate the tortuous migration behaviour. Tracks were colour coded for movement pattern, e.g. straight in red, slight turns in blue, sharp turns in yellow. Both ^msq^SPZ and ^syr^SPZ showed equal numbers of turns as compared to straight paths with the percentage turns at frame level at 88% for both samples (Fig. 4Bi). The percentage of sharp turns was somewhat decreased in ^msq^SPZ as compared to ^syr^SPZ (slight/sharp turns ^msq^SPZ: 46/42%, ^syr^SPZ: 34/54%; p<0.001, Chi-square test; Fig. 4Bi). Analysis of straightness index (Fig. 4Biii), revealed a similarly high level of tortuosity in both conditions (median straightness index ^msq^SPZ: 0.24, IQR: 0.09-0.53; ^syr^SPZ: 0.28, IQR: 0.14-0.56; p=0.14, Chi-square test). Turns were made both clockwise (CW) and counter CW (CCW), with a slight preference for CCW in the ^msq^SPZ group (CW ^msq^SPZ: 40%, ^syr^SPZ: 51%; CCW ^msq^SPZ: 60%, ^syr^SPZ: 49%; p<0.001, Chi-square test) (Fig. 4Bii). Interestingly, the turn angle of ^msq^SPZ was much more consistent, described by angular dispersion (Fig. 4Biv), as compared to ^syr^SPZ (median angular dispersion ^msq^SPZ: 0.74, IQR: 0.54-0.87; ^syr^SPZ: 0.58, IQR: 0.32-0.77; p<0.001, Chi-square test). In conclusion, both ^msq^SPZ and ^syr^SPZ travelled highly tortuous paths, whereby ^msq^SPZ showed less sharp turns, a more consistent turn angle and a predominance for the well-described preferred CCW turn angle.

#### Velocity

Sporozoite velocity fluctuated along tracks (visualized by color-coding in Fig. 5A), also in line with our earlier findings(14, 17). Plotting the average track-velocities revealed a distribution that could be described with a mixture of two normal distributions (Fig. 5B). The first peak was comparable for both administration routes and contained the slow moving ^msq^SPZ and ^syr^SPZ with a mean velocity of respectively 1.0 ± 0.4 and 0.9 ± 0.2 μm/s. The second peak, containing the highly viable and rapid ^msq^SPZ and ^syr^SPZ, differed between the administration routes at a mean of 2.4 ± 0.7 μm/s for ^msq^SPZ and 1.7 ± 0.6 μm/s for ^syr^SPZ. Thus, the rapid ^msq^SPZ moved on average 1.4-fold faster than the ^syr^SPZ.

**Figure 5.**
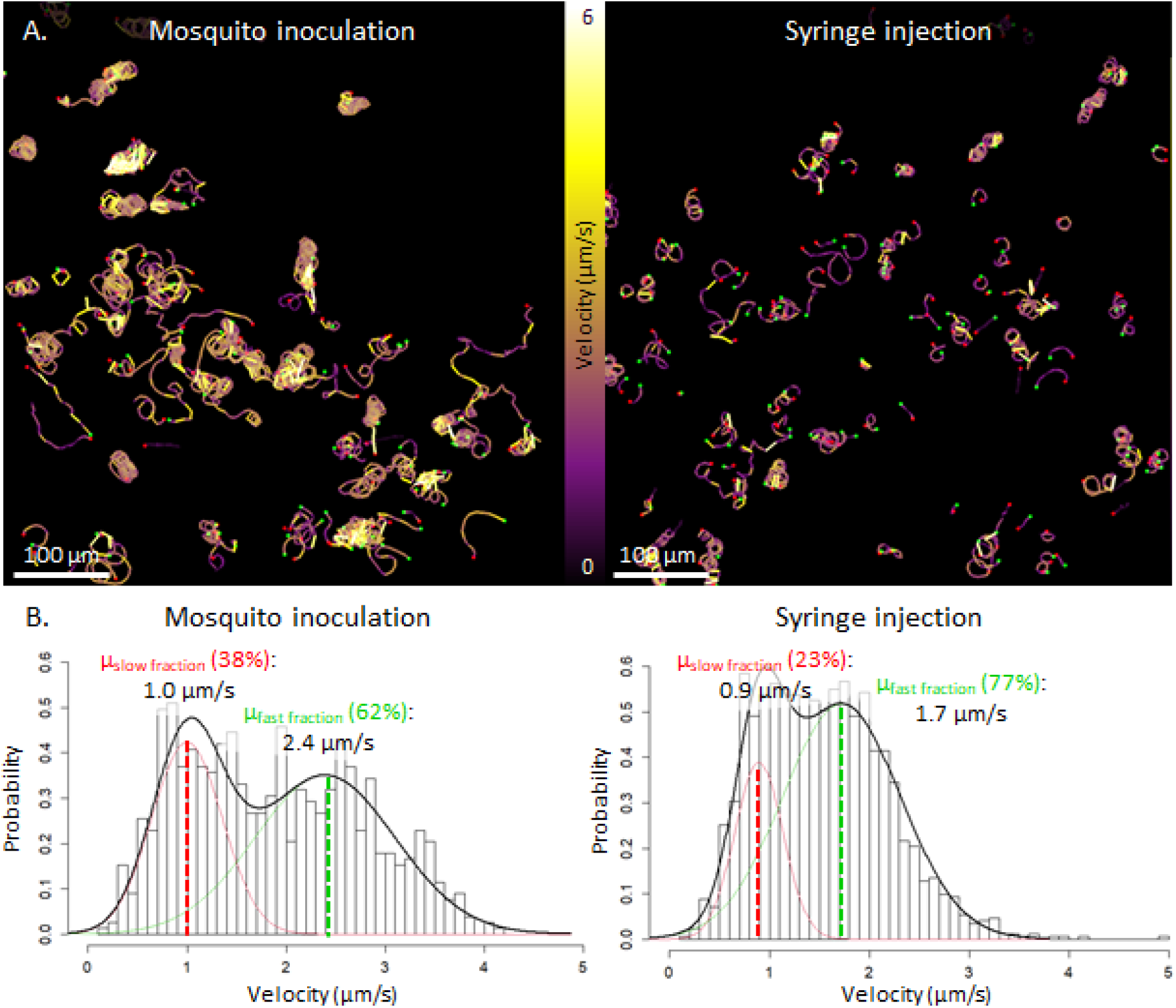
Velocity of sporozoites. (A) Overview of the inoculation site after sporozoite delivery by mosquito inoculation and the injection site after sporozoite delivery by intradermal syringe injection, shown as a map of sporozoite tracks, colour-coded for velocity (yellow sections corresponded to a high velocity, purple sections, corresponded to a lower velocity) (B) The distribution of the average track velocity including a probability density function with its mean determined using expectation-maximization based fitting of a mixture of 2 normal distributions; one describing the slow moving sporozoite fraction (depicted in red, accounting for 38% of the ^msq^SPZ and 23% of the ^syr^SPZ) and one describing the fast moving sporozoite fraction (depicted in green, accounting for 62% of the ^msq^SPZ and 77% of the ^syr^SPZ).

### Interplay between motility parameters

To obtain a multidimensional view of sporozoite migration, we explored the relationship between tortuosity parameters. Based on high and low straightness index (SI) and angular dispersion (AD), the sporozoites tracks could be divided into four typical movement patterns: short erratic tracks (low AD, high SI), short straight tracks (high AD, high SI), consistently turning tracks (high AD, low SI) and erratically turning tracks (low AD, low SI) (Fig. 6A). Representative examples of tracks from these groups are shown in Fig. 6B. The majority of the tracks (49%) were classified as consistently turning (49%), which typically is the “default” movement pattern which sporozoites display *in vitro* (Fig. 6A).

**Figure 6.**
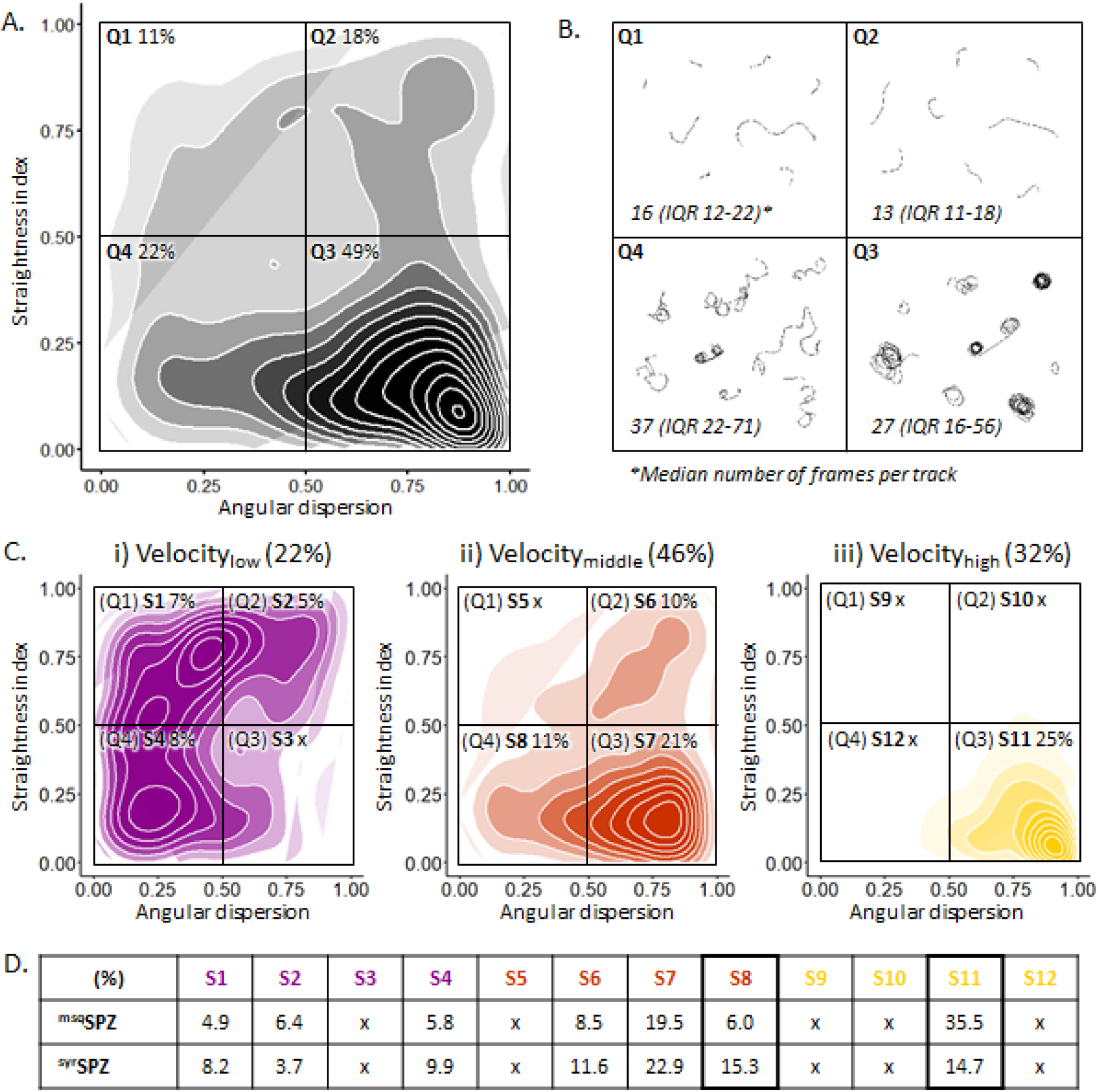
Interplay between motility parameters. (A) Density plot of all sporozoite tracks (including both ^msq^SPZ (n=778) and ^syr^SPZ (n=778) in a matrix of angular dispersion and the straightness index. (B) Representative example movement patterns associated with every quarter of the density plot. Group Q1: low angular dispersion, high straightness index (short, erratic tracks); Q2: high angular dispersion, high straightness index (short, straight tracks); Q3: high angular dispersion, low straightness index (consistently turning tracks); Q4: low angular dispersion, low straightness index (erratically turning tracks). (C) Density plot of all low (i), intermediate (ii) and high (iii) velocity tracks on the same matrix of angular dispersion and straightness index, resulting in twelve subsets (S1-12). Velocity categories are defined as: slow <1 μm/s (depicted in purple), intermediate 1-2 μm/s (depicted in orange) and fast >2 μm/s (depicted in yellow). Percentage of the overall number of sporozoite tracks in every quarter of the density plots are given. (D) Comparison of the distribution (as a percentage of total tracks) for ^msq^SPZ and ^syr^SPZ tracks across subsets S1-12, with significant differences between S8 and S11 (p<0.001, Chi-square test).

We subsequently investigated the relationship between the four movement patterns and velocity. We found that consistently turning tracks were generally rapid sporozoites (median 2.0 μm/s, IQR 1.5-2.6), whereas erratically turning sporozoites were considerably slower (median: 1.2 μm/s, IQR 0.9-1.7; p<0.001, Mann-Whitney U test). Interestingly, these consistently turning rapid sporozoites were overrepresented within the ^msq^SPZ group as compared to the ^syr^SPZ group (2.4-fold difference), which was offset by a reciprocal increase in slower erratically turning sporozoites within the ^syr^SPZ group (2.5-fold difference) (p<0.001, Chi-square test) (Fig. 6D).

Lastly, we determined if the distribution of the sporozoites as reflected by the nearest neighbour distance (NND) influenced the tortuosity and velocity of their tracks. In general, the average NND of ^msq^SPZ was larger compared to ^syr^SPZ as was observed by the analysis of the individual parameters (Fig. 2, Sup. Fig. 4). However, this difference was consistent among all different movement patterns (p=0.52; interaction term ^msq/syr^SPZ * subsets, univariate general linear model), which suggested that the movement pattern or velocity of sporozoites was not dependent on their interindividual distance.

Taken together, the ^syr^SPZ group contained more sporozoites that exhibited erratic movement at a slower speed, whereas the ^msq^SPZ group contained more sporozoites that circled consistently at high speed.

## DISCUSSION

Using confocal microscopy and dedicated sporozoite imaging software, we visualized sporozoites deposited by mosquito inoculation or syringe injection and assessed quantitative differences. We found that delivery by syringe injection decreases infectivity of sporozoites by three-fold as compared to mosquito inoculation, which is related to 1) a clustered distribution of ^syr^SPZ through the skin, 2) a lack of hematomas which are typically induced by mosquito bites, 3) enlarged interstitial space due to syringe injection pressure, 4) slower and more erratic migration patterns of ^syr^SPZ as compared to ^msq^SPZ. Each of these parameters could impact the efficiency of sporozoite migration, blood vessel invasion and ultimate liver infectivity and thus provide important insights how to critically improve delivery of sporozoite-based vaccines.

The dispersed distribution of ^msq^SPZ and the fact that a substantial proportion of ^msq^SPZ are deposited close to hematomas created by mosquito probing(23, 24) provides them with better odds of finding blood stream access as compared to ^syr^SPZ. This is in line with the consistent circular motility of ^msq^SPZ, which has previously been associated with increased blood vessel engagement(13, 16). However, in previous publications this engagement was generally related to deceleration, while in our study consistent circular motility was related to high velocities(13, 16). Conversely, the slower erratic movement of ^syr^SPZ is most likely caused by the altered physical space as a consequence of liquid injection pressure. Sporozoite movement is strongly guided by their three-dimensional environment; without any confinement, sporozoites *in vitro* display a continuous, preferential counter-clockwise movement pattern(14, 25), while *in vivo*, skin structure redirects sporozoites to display much more complex patterns(15, 17). The role of physical confinement has been further supported by experiments whereby micropatterned *in vitro* environments were created that could induce specific movement patterns of sporozoites(26). We clearly found that the liquid which was co-injected with the ^syr^SPZ widened the interstitial space, which allows ^syr^SPZ to display erratic movement patterns.

Despite the fact that mouse skin does not fully replicate human skin with regards to skin thickness (mouse skin <1 mm vs human skin >2 mm(27)), as underlined by ^msq^SPZ deposition in the mouse peritoneum (length of mosquito proboscis: 1.5-2.0 mm(28)), the remarkable differences between the ^msq^SPZ and ^syr^SPZ dermal sites provide important clues how to critically improve intradermal syringe injections of attenuated sporozoite vaccines. Particularly, a microneedle (patch) or tattoo device may be useful to not only create the relevant sporozoite dispersion, but also to significantly decrease injection volume and tissue pressure(29–32). In addition, laser-induced vascular damage can potentially mirror the hematomas induced by mosquito probing and enhance the blood vessel entrance of ^syr^SPZ(33). Recently, this concept was successfully applied to increase parasite loads in the liver after intradermal syringe injection(34).

Importantly, our study demonstrates that state-of-the-art imaging (analysis) techniques can provide valuable quantitative assessments of parameters affecting sporozoite migration. Our *ex vivo* set-up combined with spinning-disk confocal microscopy and sporozoite tracking software enabled 1) the visualization of (the sporozoite distribution throughout) the whole inoculation and injection site (up to 100 mm^2^), whereas up to now only one field of view (<0.5 mm^2^) was visualized during *in vivo* live imaging(13, 16, 35), 2) the visualization of morphological tissue deformation as a result of pressure of fluid injection, previously acknowledged as an important parameter regarding transdermal drug delivery(36–38), but not yet visualized at a micron level resolution and 3) a multidimensional analysis of sporozoite motility unveiling an remarkable interplay between motility parameters which were up to now only studied independently(15–17). Further research is needed to study the role of other potential contributors not accounted for in this imaging study, such as their pre-processing (manual extraction from salivary glands in culture medium instead of saliva(14, 39, 40)). Our quantitative assessment of parameters affecting sporozoite migration both indicates that engineering solutions should be explored that can better mimic mosquito inoculation as well as provides a readout needed to assess the potential of the suggested engineering solutions.

## CONCLUSION

In conclusion, detailed microscopic imaging of the dermal site appearance and migration patterns of sporozoites revealed important quantitative differences between sporozoite administration via mosquito inoculation vs intradermal syringe injection. These findings open new avenues for intradermal delivery of attenuated sporozoite vaccines with enhanced efficacy.

## ACKNOWLEDGEMENTS

We would like to thank ing. J. Ramesar and dr. C.J. Janse for support with the mosquito infection. We also thank the light microscopy facility team of the LUMC for their support during image acquisition and analysis.

## SUPPLEMENTARY DATA

**Supplementary Figure 1.**
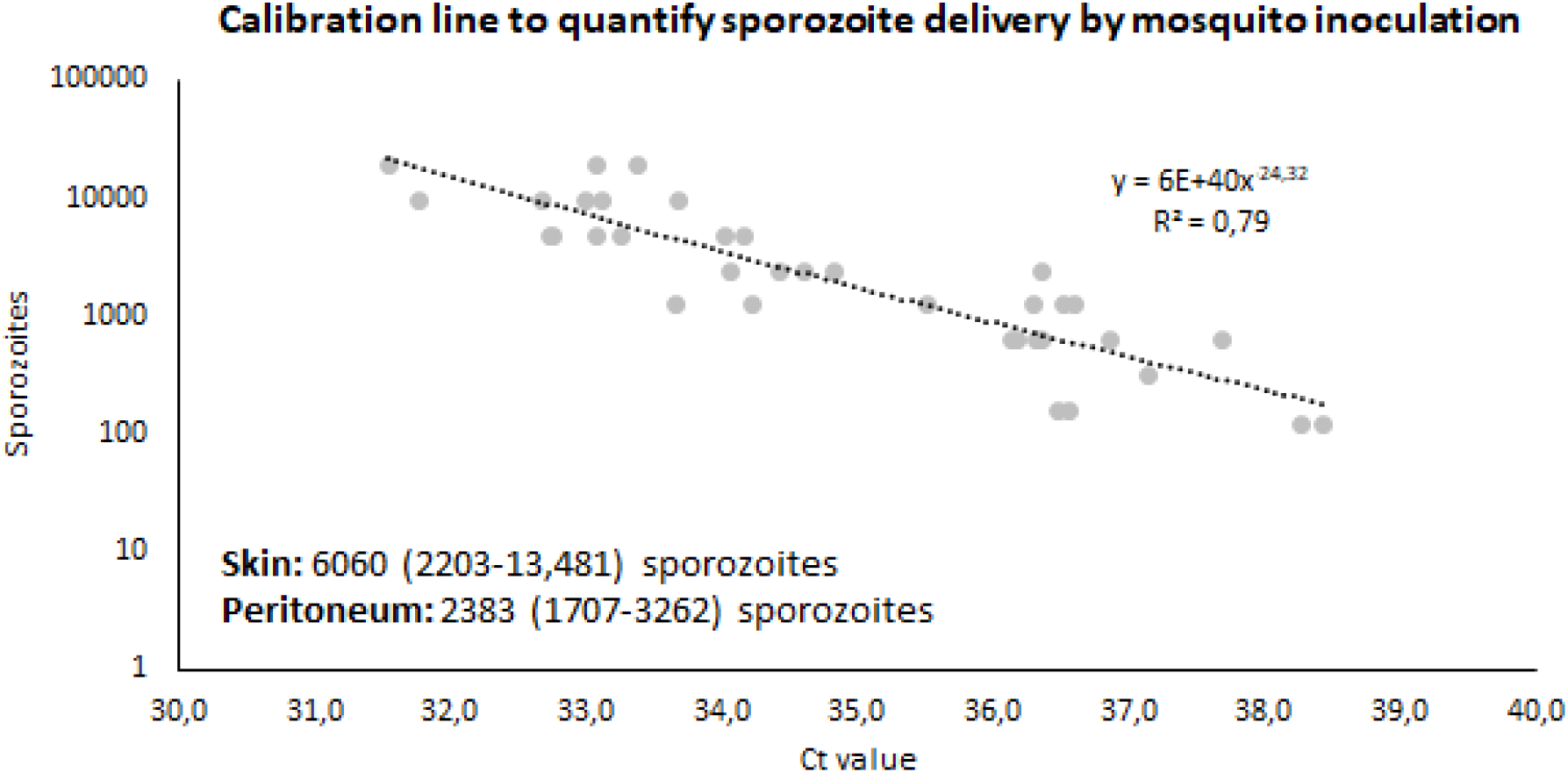
Sporozoite delivery quantification. A calibration curve was generated based on a syringe injected concentration range of sporozoites in skin (n=3 in duplo) to estimate the number of sporozoites delivered by 30 mosquito bites.

**Supplementary Figure 2.**
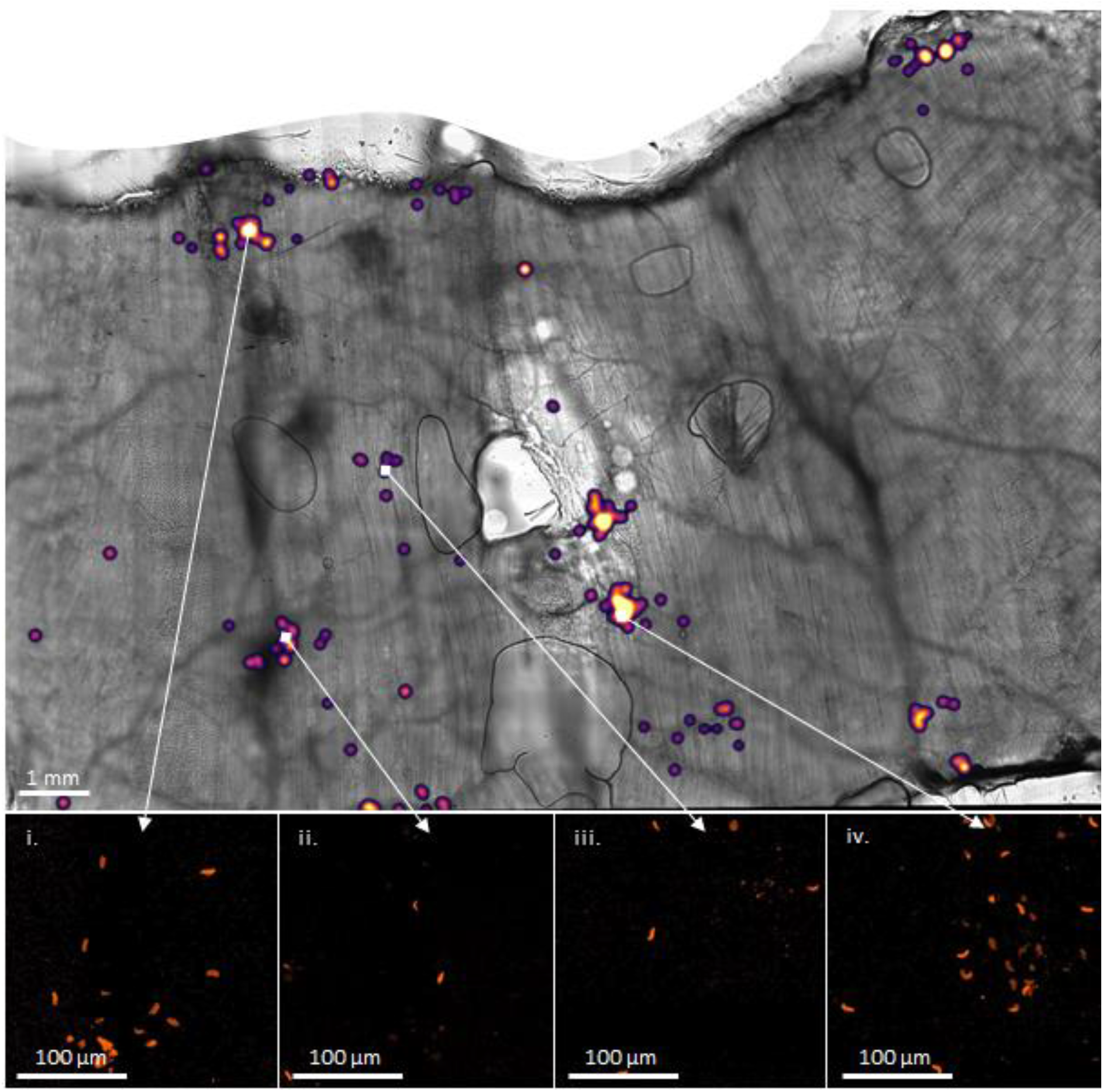
Peritoneum after mosquito bites. Overview of peritoneum after sporozoite delivery by mosquito inoculation shown as an overlay of a brightfield image and the sporozoite distribution accompanied by zoom-in images showing the individual sporozoites (i-iv).

**Supplementary Figure 3.**
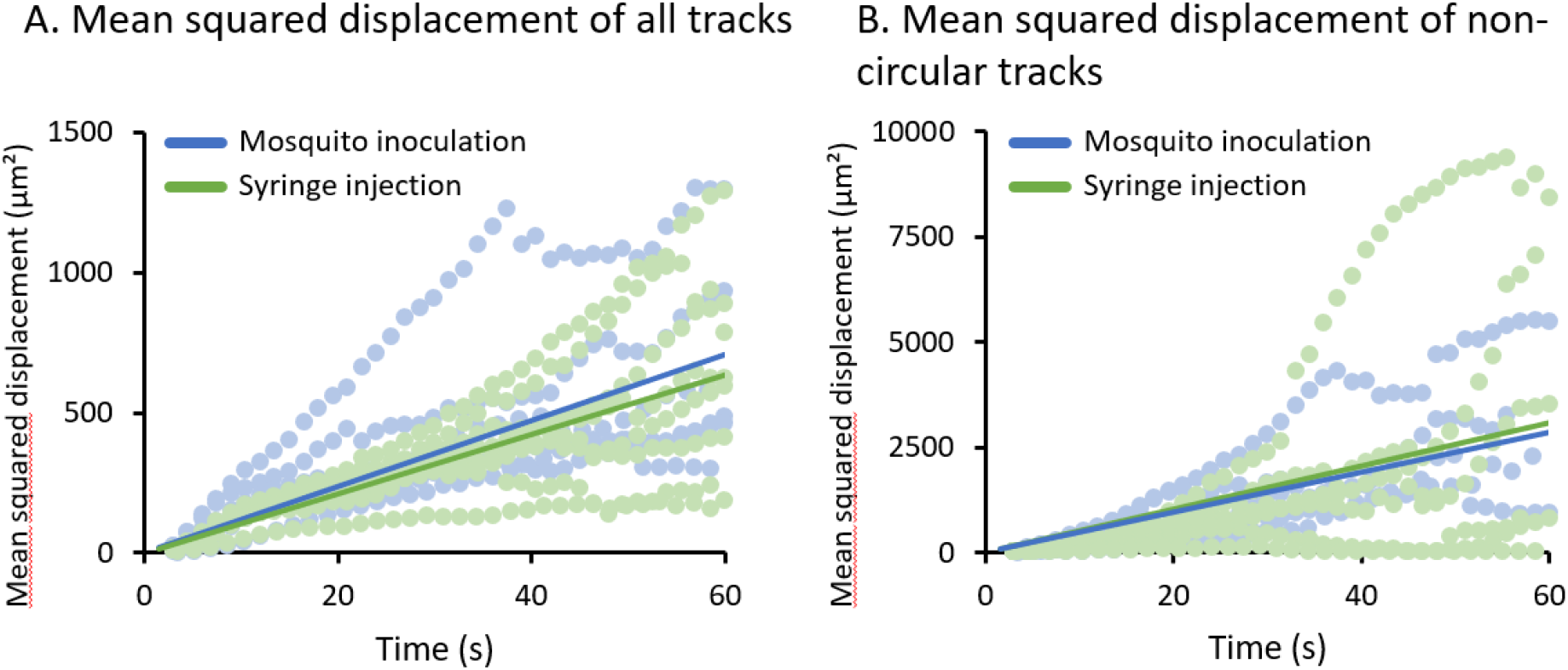
Mean squared displacement. A-B) The mean squared displacement of the sporozoites after administration by mosquito inoculation or syringe injection is plotted. In A) all tracks are included, in B) the circular tracks are excluded (90% of mosquito inoculation frames, 95% of syringe injection frames). The mean squared displacement of the individual samples is plotted as a dotted line, a linear trendline is plotted as a solid line.

**Supplementary Figure 4.**
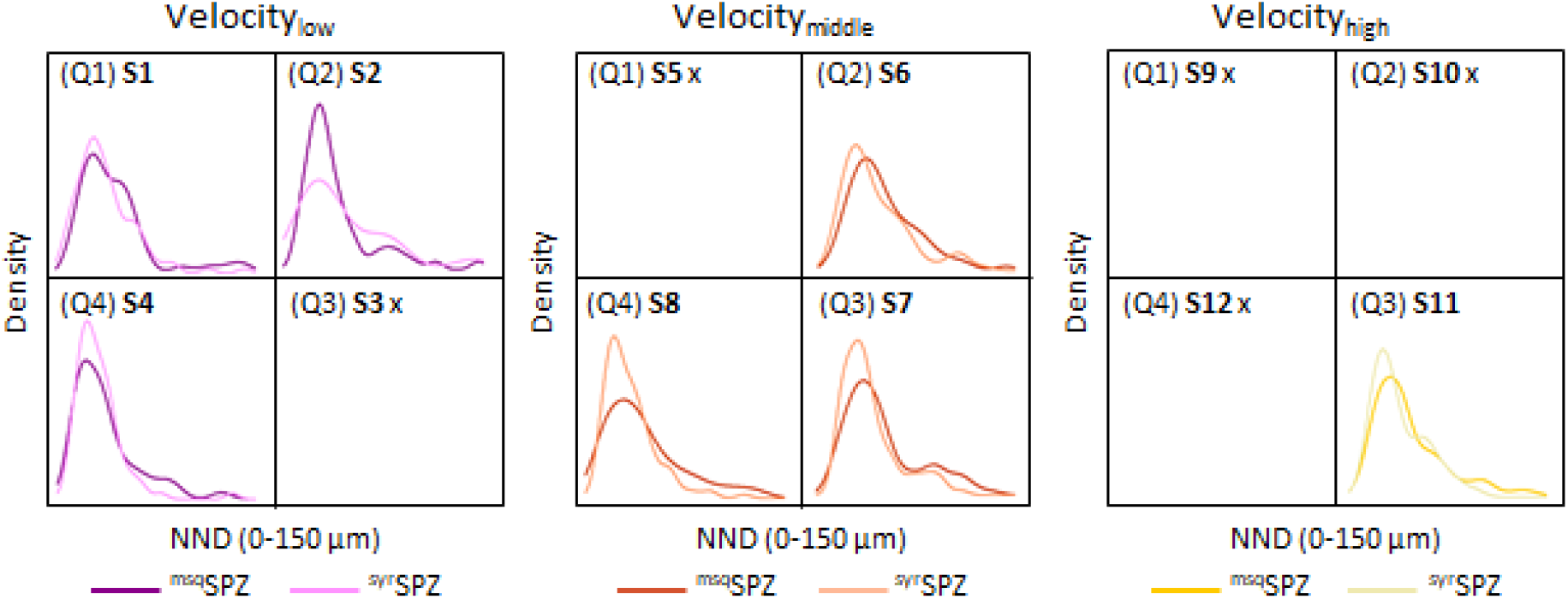
Relation between motility parameters and nearest neighbour distance. The nearest neighbour distance (NND) is plotted for the tracks split into 12 subsets (S1-12) based on four specific movement patterns (Q1-4, see Fig. 6) and three velocity categories (1: <0.1 μm/s, depicted in purple; 2: 1-2 μm/s, depicted in orange; 3: >2 μm/s, depicted in yellow). The NND of the ^msq^SPZ is depicted in the darker colour and the NND of the ^syr^SPZ in depicted in the lighter colour.

